# A New Resource for Genomics and Precision Health Information and Publications on the Investigation and Control of COVID-19 and other Coronaviruses

**DOI:** 10.1101/2020.04.21.050922

**Authors:** Wei Yu, Marta Gwinn, Muin J. Khoury

## Abstract

**Summary:** We developed a new online database that contains the most updated published scientific literature, online news and reports, CDC and National Institutes of Health (NIH) resources. The tool captures emerging discoveries and applications of genomics, molecular, and other precision medicine and precision public health tools in the investigation and control of coronavirus diseases, including COVID-19, MERS-CoV, and SARS.

**Availability:** Coronavirus Disease Portal (CDP) can be freely accessed via https://phgkb.cdc.gov/PHGKB/coVInfoStartPage.action.

## 1 INTRODUCTION

Advances in genomics and precision health continue to influence the evolution of medicine and public health (Feero, 2017). In particular, the emerging use of pathogen genomics has ushered in a new era of precision public health (Armstrong, 2019). The COVID-19 pandemic has catalyzed the scientific research community to investigate the biology, epidemiology, and clinical manifestations of infection with this novel virus, SARS-CoV-2, as well as potential therapeutic and preventive interventions. .Hundreds of articles have already been published in the peer reviewed literature and nearly two thousand more in preprint servers.

How are new tools of genomics and precision health and technologies being used to investigate and control COVID-19? As part of an effort to help translate new technologies into public health benefit, our office at the Centers for Disease Control and Prevention (CDC) has created a freely accessible, web-based application called Coronavirus Disease Portal (CDP) in Genomics and Precision Health,^1^ a component of the CDC Public Health Genomics and Precision Health Knowledge Base (PHGKB).^2^ PHGKB was launched in 2016 as a tool for researchers, clinicians and public health professionals interested in the translation of research on human and pathogen genomics, big data and machine learning, and other precision health technologies into population health benefits. PHGKB has been described in detail elsewhere and features a suite of curated and continuously updated databases (Yu et al, 2016).

## 2 IMPLEMENTATION

CDP is a web-based application based on J2EE technology^3^ with Java open-source frameworks including Hibernate^4^ and Strut.^5^ As one component of the PHGBKB system, CDP has been integrated into the overall architecture of PHGKB, described previously (Yu et al, 2007a). PubMed records are one of main data resources and are e indexed with MeSH terminology;^6^; the MeSH tree hierarchies and the Unified Medical Language System ^7^ metathesaurus enhance the search capacity of the system.

Data in the database are collected mainly through automatic retrieval from PubMed and manual curation by domain experts at CDC. CDP automatically retrieves scientific literature in genomics and precision health daily from PubMed with two specifically designed queries: Genomics Precision Health and Non-Genomics Precision Health (Appendix). An automatic PubMed script using NCBI Eutils^8^ is used to retrieve records automatically, based on the two PubMed queries, and to identify records related to COVID-19. Online news and reports related to coronavirus are selected by CDC staff from our weekly horizon scan for Genomics Health Impact Update^9^ and CDC Advanced Molecular Detection (AMD) Clips.^10^ A web-based curation pipeline allows CDC domain experts to select and curate important news, reports, and articles related to COVID-19 and other coronaviruses. A text mining technique (Yu et al, 2007b) is used to identify and standardize the country information associated with the authors in PubMed records. All data selection processes are performed daily.

## 3 DESCRIPTION OF FEATURES

The new portal is an online, searchable database of published scientific literature, CDC and NIH resources, and other information. It is designed to capture emerging discoveries and applications of genomics, molecular, and other precision health technologies in the investigation and control of coronaviruses, including MERS-CoV and SARS, with a particular focus on SARS-CoV-2. The records selected are further classified into categories: SARS-CoV-2 versus other coronaviruses, and genomic versus non-genomic. The CDP links to other curated contents of PHGKB and selected external resources. CDP is designed as a user-friendly interface (Figure 1) with the following functionalities: 1) Free text search can be performed as an option in either the entire PHGKB dataset or limited by subcategories (e.g., COVID-19 only, genomics only); 2) Search results can be further filtered by publication year, journal, country of the authors; 3) A graph of the number of publications by year is generated dynamically for the search results; 4) Search results can be downloaded in tab-delimited text format; 5) User can sign up for a daily email alert that contains any new updated content. As of April 21, 2020, CDP contained 6123 records, including 5610 PubMed abstracts by authors from 111 countries, published since 2000 in 910 journals. Among these are 465 publications related to COVID-19, most published by Chinese researchers in 2020. CDP also includes 154 publications on preprint servers (bioRxiv.org and medRxiv.org), as well as 513 online news stories and reports.

**Figure 1:**
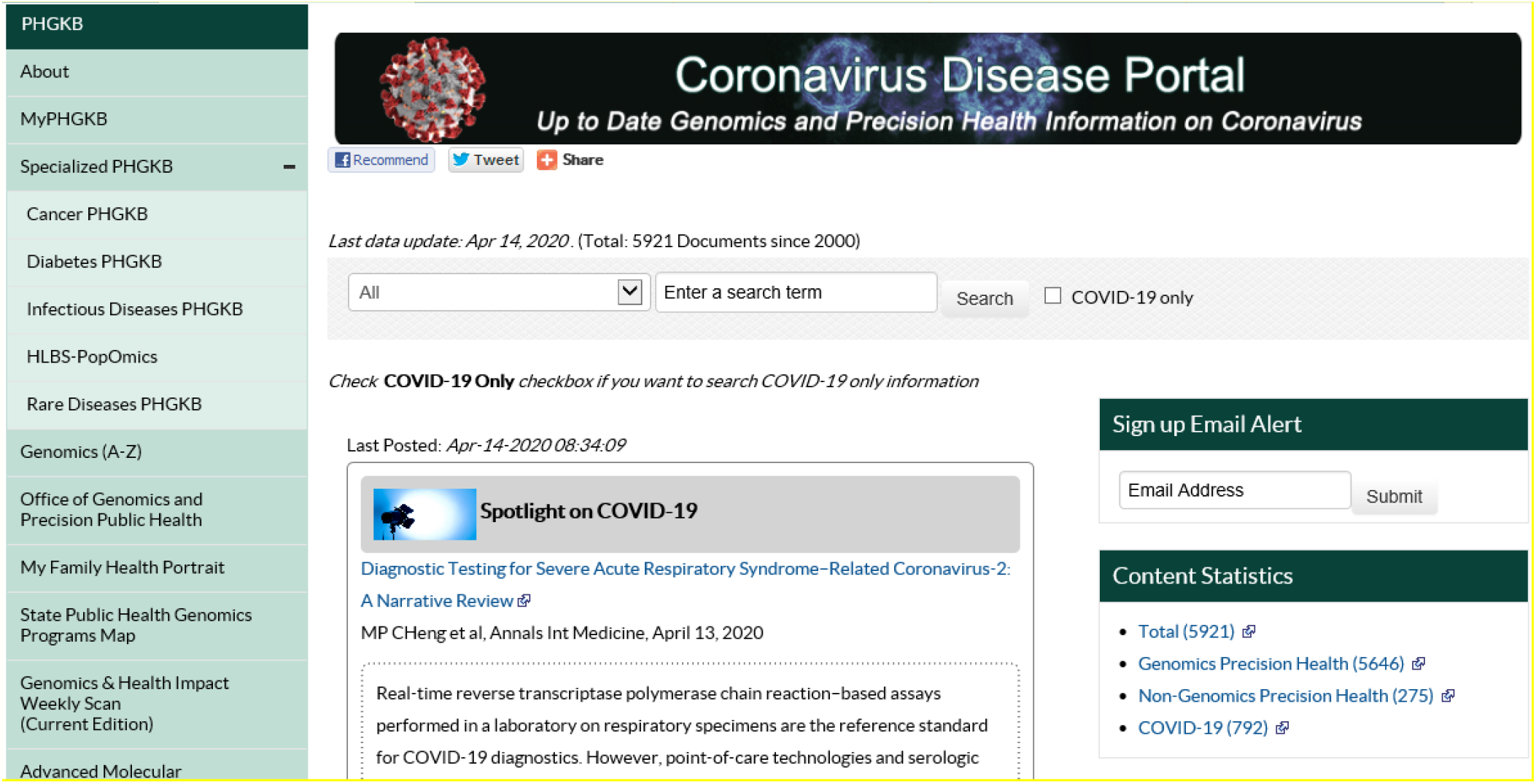
The home page of the Coronavirus Disease Portal in Genomics and Precision Health

## 4 EXAMPLES OF THE USAGE

Using CDP, users can dynamically create quick snapshots of research on coronavirus or COVID-19 based on any combination of variables available in the dataset including publication type (PubMed vs other), publication year, publication journal, country of authors, and genomic vs. non-genomic.

Examples of possible analyses using CDP data as of April 21, 2020 include:
 
1. Trend in number of PubMed publications by year on all coronaviruses and COVID-19 (Figure 2). This analysis shows three peaks in the number of scientific publications, reflecting three major epidemics of coronavirus (SARS, MERS, and COVID-19).
2. The top 10 countries reporting coronavirus research in genomics and precision health (Table 1). China ranks highest, followed by United States.
3. The top 10 journals reporting coronavirus research in genomics and precision health (Table 2). The Journal of Virology ranks highest.

**Figure 2.**
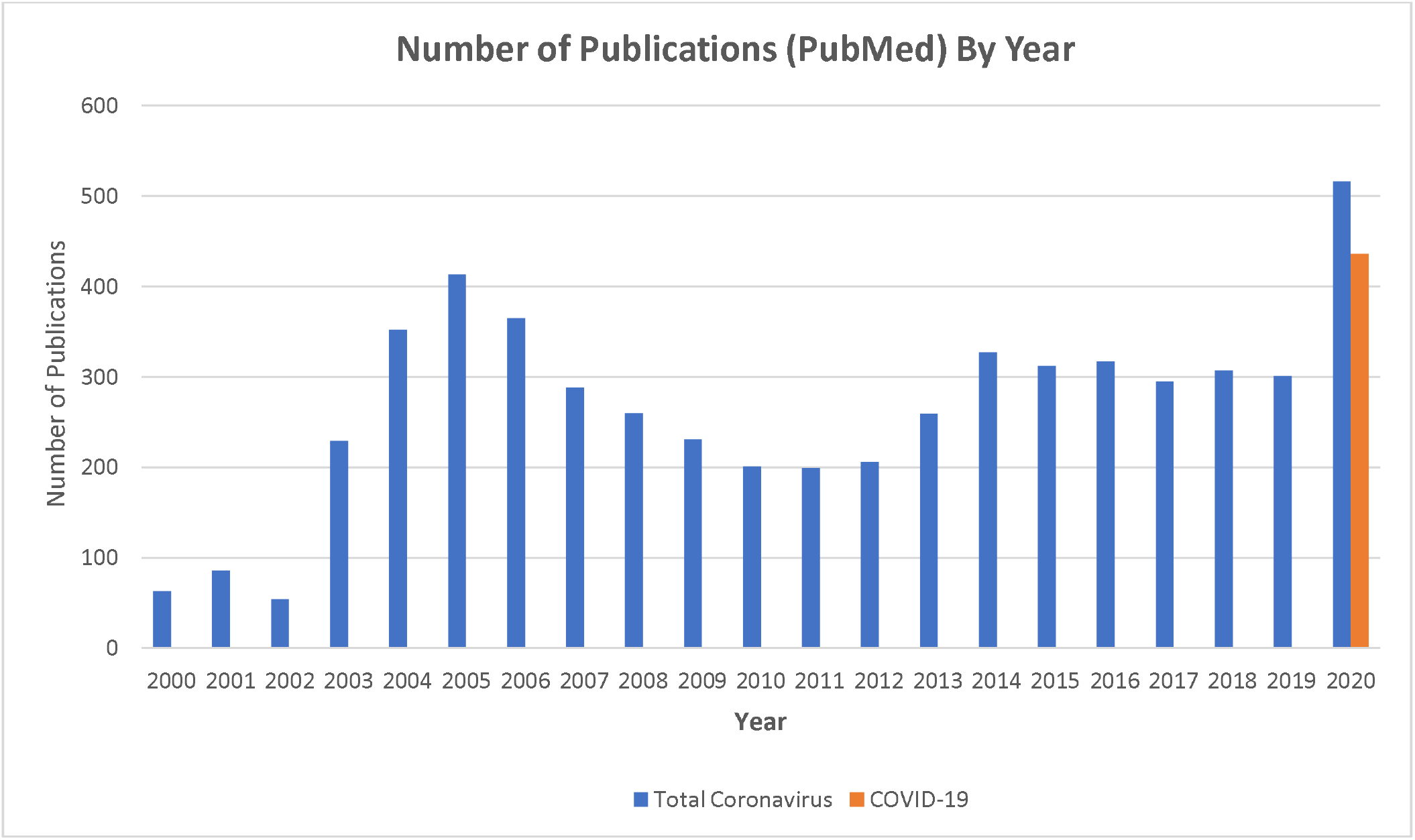
Trend of Number of Publications (PubMed) over Year in Coronavirus and COVID-19.

**Table 1.** Top 10 Countries in Publications in Genomics Precision Health Research.

| **Country** | **Number of Publication**<br>**(PubMed)** |
| --- | --- |
| China | 100 |
| United States | 41 |
| Italy | 22 |
| United Kingdom | 12 |
| India | 8 |
| Australia | 8 |
| Brazil | 7 |
| France | 7 |
| South Korea | 7 |
| Singapore | 7 |

**Table 2.** Top 10 Journals in Publications in Coronavirus Research.

| **Journal** | **Number of Publications (PubMed)** |
| --- | --- |
| Journal of virology | 626 |
| Virology | 196 |
| The Journal of general virology | 118 |
| PloS one | 116 |
| Virus research | 115 |
| Emerging infectious diseases | 107 |
| Journal of medical virology | 93 |
| Archives of virology | 91 |
| Journal of virological methods | 90 |
| Viruses | 86 |

## 5 DISCUSSION

COVID-19 is the biggest public health challenge of our times. The rapid emergence of a novel coronavirus has accelerated the application of genomics and big data tools and technologies to its investigation and control. Our new portal captures diverse examples of such applications, including the use of whole genome sequencing to track the virus’s origin and spread, including detailed geographic information at the global, country, and local levels; the use of smartphone based tracking and control; and rapid characterization of risk factors related to severe disease such as age and underlying medical conditions. The role of host genomic factors is beginning to be explored. Machine learning and data science are contributing to diagnosis and prediction of complications. Emerging digital technologies could augment traditional public-health strategies for tackling COVID-19 including public health surveillance, detection, and control, and mitigation of impact on healthcare delivery.

The information accessible CDP represents about 15% of the total scientific literature and resources on COVID-19, selected to focus on the use of genomics and other precision health tools in scientific studies of COVID-19. We are continuously updating this resource as more data become available. We hope that CDP can serve as a scientific information portal to facilitate coronavirus research in genomics and precision public health.

## Foot Notes

1 https://phgkb.cdc.gov/PHGKB/coVInfoStartPage.action

2 https://phgkb.cdc.gov/

3 http://java.sun.com/javaee/

4 http://www.hibernate.org/

5 http://struts.apache.org/

6 https://www.ncbi.nlm.nih.gov/mesh

7 https://www.nlm.nih.gov/research/umls/index.html

8 https://www.ncbi.nlm.nih.gov/books/NBK25501/

9 https://phgkb.cdc.gov/PHGKB/translationClip.action?action=home

10 https://phgkb.cdc.gov/PHGKB/amdClip.action?action=home

## References

Armstrong GL, MacCannell DR, Taylor J, Carleton HA, Neuhaus EB, Bradbury RS, Posey JE, Gwinn M. Pathogen genomics in public health. N Engl J Med. 2019 Dec 26;381(26):2569–2580.

Feero WG. An introduction to genomics and precision health JAMA 2017;317(18):1842–1843.

Yu, W., Gwinn M, Dotson WD, et al. A knowledge base for tracking the impact of genomics on population health Genet Med. 2016 Dec;18(12):1312–1314

Yu W, Yesupriya A, Wulf A, et al. An open source infrastructure for managing knowledge and finding potential collaborators in a domain-specific subset of PubMed, with an example from human genome epidemiology. BMC Bioinformatics. 2007 Nov 9;8:436.

Yu W, Yesupriya A, Wulf A, et al. An automatic method to generate domain-specific investigator networks using PubMed abstracts. BMC Med Inform Decis Mak 2007 Jun 20;7(1):17.

