## Appendix for "A New Resource for Genomics and Precision Health Information and Publications on the Investigation and Control of COVID-19 and other Coronaviruses"

PubMed queries:

**Coronavirus and Genomics Precision Health**

(((((((((\"coronavirus\"[MeSH Terms] OR \"coronavirus\"[All Fields]) OR \"coronaviruses\"[All Fields]) OR \"MERS-Cov\"[All Fields]) OR \"SARS-CoV\"[All Fields]) OR \"SARS-CoV-2\"[All Fields]) OR \"COVID-19\"[All Fields]) OR \"2019-nCoV\"[All Fields]) OR \"Severe Acute Respiratory Syndrome\"[All Fields]) OR \"Middle East Respiratory Syndrome\"[All Fields]) AND ((((((((((((((((((((((\"bioinformatical\"[All Fields] OR \"bioinformatically\"[All Fields]) OR \"computational biology\"[MeSH Terms]) OR (\"computational\"[All Fields] AND \"biology\"[All Fields])) OR \"computational biology\"[All Fields]) OR \"bioinformatic\"[All Fields]) OR \"bioinformatics\"[All Fields])) OR ((((((((((((\"genetic therapy\"[MeSH Terms] OR (\"genetic\"[All Fields] AND \"therapy\"[All Fields])) OR \"genetic therapy\"[All Fields]) OR \"genetic\"[All Fields]) OR \"genetical\"[All Fields]) OR \"genetically\"[All Fields]) OR \"genetics\"[MeSH Subheading]) OR \"genetics\"[All Fields]) OR \"genetics\"[MeSH Terms]) OR \"genet\"[All Fields]) OR \"genets\"[All Fields])) OR (((((((\"genome\"[MeSH Terms] OR \"genome\"[All Fields]) OR \"genomes\"[All Fields]) OR \"genome s\"[All Fields]) OR \"genomically\"[All Fields]) OR \"genomics\"[MeSH Terms]) OR \"genomics\"[All Fields]) OR \"genomic\"[All Fields])) OR ((\"genes\"[MeSH Terms] OR \"genes\"[All Fields]) OR \"gene\"[All Fields])) OR ((((((((((\"mutate\"[All Fields] OR \"mutated\"[All Fields]) OR \"mutates\"[All Fields]) OR \"mutating\"[All Fields]) OR \"mutation\"[MeSH Terms]) OR \"mutation\"[All Fields]) OR \"mutations\"[All Fields]) OR \"mutation s\"[All Fields]) OR \"mutational\"[All Fields]) OR \"mutator\"[All Fields]) OR \"mutators\"[All Fields])) OR (((((((((((\"genotype\"[MeSH Terms] OR \"genotype\"[All Fields]) OR \"genotypes\"[All Fields]) OR \"genotypic\"[All Fields]) OR \"genotype s\"[All Fields]) OR \"genotyped\"[All Fields]) OR \"genotyper\"[All Fields]) OR \"genotypical\"[All Fields]) OR \"genotypically\"[All Fields]) OR \"genotyping\"[All Fields]) OR \"genotypings\"[All Fields]) OR \"genotypization\"[All Fields])) OR (((((((\"genome\"[MeSH Terms] OR \"genome\"[All Fields]) OR \"genomes\"[All Fields]) OR \"genome s\"[All Fields]) OR \"genomically\"[All Fields]) OR \"genomics\"[MeSH Terms]) OR \"genomics\"[All Fields]) OR \"genomic\"[All Fields])) OR (((((((((((\"genotype\"[MeSH Terms] OR \"genotype\"[All Fields]) OR \"genotypes\"[All Fields]) OR \"genotypic\"[All Fields]) OR \"genotype s\"[All Fields]) OR \"genotyped\"[All Fields]) OR \"genotyper\"[All Fields]) OR \"genotypical\"[All Fields]) OR \"genotypically\"[All Fields]) OR \"genotyping\"[All Fields]) OR \"genotypings\"[All Fields]) OR \"genotypization\"[All Fields])) OR (((((((\"polymorphic\"[All Fields] OR \"polymorphics\"[All Fields]) OR \"polymorphism s\"[All Fields]) OR \"polymorphism, genetic\"[MeSH Terms]) OR (\"polymorphism\"[All Fields] AND \"genetic\"[All Fields])) OR \"genetic polymorphism\"[All Fields]) OR \"polymorphism\"[All Fields]) OR \"polymorphisms\"[All Fields])) OR ((((((((((((\"allel\"[All Fields] OR \"allele s\"[All Fields]) OR \"alleleic\"[All Fields]) OR \"alleles\"[MeSH Terms]) OR \"alleles\"[All Fields]) OR \"allele\"[All Fields]) OR \"allelic\"[All Fields]) OR \"allelically\"[All Fields]) OR \"allelism\"[All Fields]) OR \"allelisms\"[All Fields]) OR \"allelle\"[All Fields]) OR \"allelles\"[All Fields]) OR \"allels\"[All Fields])) OR ((\"variant\"[All Fields] OR \"variant s\"[All Fields]) OR \"variants\"[All Fields])) OR ((((((((((((\"allel\"[All Fields] OR \"allele s\"[All Fields]) OR \"alleleic\"[All Fields]) OR \"alleles\"[MeSH Terms]) OR \"alleles\"[All Fields]) OR \"allele\"[All Fields]) OR \"allelic\"[All Fields]) OR \"allelically\"[All Fields]) OR \"allelism\"[All Fields]) OR \"allelisms\"[All Fields]) OR \"allelle\"[All Fields]) OR \"allelles\"[All Fields]) OR \"allels\"[All Fields])) OR (((\"heterozygote\"[MeSH Terms] OR \"heterozygote\"[All Fields]) OR \"heterozygotes\"[All Fields]) OR \"heterozygotic\"[All Fields])) OR (((((\"haplotyped\"[All Fields] OR \"haplotypes\"[MeSH Terms]) OR \"haplotypes\"[All Fields]) OR \"haplotype\"[All Fields]) OR \"haplotypic\"[All Fields]) OR \"haplotyping\"[All Fields])) OR (((\"genome-wide association study\"[MeSH Terms] OR ((\"genome wide\"[All Fields] AND \"association\"[All Fields]) AND \"study\"[All Fields])) OR \"genome wide association study\"[All Fields]) OR \"gwas\"[All Fields])) OR \"genome wide association\"[All Fields]) OR \"hereditary\"[All Fields])

**Coronavirus and Non-Genomics Precision Health**

(((((((((\"coronavirus\"[MeSH Terms] OR \"coronavirus\"[All Fields]) OR \"coronaviruses\"[All Fields]) OR \"MERS-Cov\"[All Fields]) OR \"SARS-CoV\"[All Fields]) OR \"SARS-CoV-2\"[All Fields]) OR \"COVID-19\"[All Fields]) OR \"2019-nCoV\"[All Fields]) OR \"Severe Acute Respiratory Syndrome\"[All Fields]) OR \"Middle East Respiratory Syndrome\"[All Fields]) AND (((((((\"Big data\"[All Fields] OR \"Data science\"[All Fields]) OR \"Machine Learning\"[All Fields]) OR \"Artificial Intelligence\"[All Fields]) OR \"Predictive Analytics\"[All Fields]) OR \"Digital Health\"[All Fields]) OR \"Deep Learning\"[All Fields]) OR \"Neural Networks\"[All Fields])
